## Supplementary Materials for "Intrasexual cuticular hydrocarbon dimorphism in a wasp sheds light on hydrocarbon biosynthesis genes in Hymenoptera"

### Supplementary Methods

#### *Trap nests*

*Odynerus spinipes* trap nests consisted of 20 cm long plastic pipes with a clear diameter of 10 cm filled with eolian silt deposit. The eolian silt deposit contained pre-built 7 cm long channels with a diameter of 0.6 cm each. The trap nests were mounted on posts at a height between 5 and 100 cm from the ground in Büchelberg, Germany (49.027985, 8.164801) in February 2015. The prepupae were removed from the trap nests in March 2016 and stored in separate transparent gelatin capsules (empty hard gelatin capsules size 3, LUTOR trading & distribution, Cologne, Deutschland and Birmingham, United Kingdom). Each gelatin capsule was pierced with an insect needle to improve gas exchange. The prepupae were stored at 4 °C within a box containing a wet tissue (changed every two days). Synchronized development was triggered end of April 2016 by increasing the ambient temperature to 23 °C. This resulted in most males hatching one week and most females hatching two weeks later.

*O. spinipes* females used for transcriptome sequencing were collected from trap nests placed in Bad Muskau (51.548200, 14.714900; in 2017), Büchelberg (49.027985, 8.164801; in 2015 and 2017), and Eichenzell (50.495927, 9.704589; in 2016), and treated as described above.

#### *Gas chromatography and mass spectrometry*

CHC extracts of *Odynerus spinipes* samples kept in the laboratory (from Büchelberg [2015 and 2016] and from Eichenzell [2016]) as well as of the three wasps sampled at the field site Büchelberg in 2016 were processed with an Agilent 6890 gas chromatograph (GC) coupled to an Agilent 5975 mass selective detector (MS, Agilent Technologies, Waldbronn, Germany). The GC was operated in splitless injection mode and fitted with a DB-5 Fused Silica capillary column (30 m × 0.25 mm ID; film thickness: 0.25 µm; J&W Scientific, Folsom, United States). CHC extracts of wasps sampled in 2017 in the field at the field site Büchelberg were processed with HP series 6890 gas chromatograph (Hewlett Packard, Palo Alto, California, USA), equipped with a DB-5 column (30 m x 0.25 mm film thickness: 0.25 µm; J&W Scientific, Folsom, USA), coupled to a HP series 5973 quadrupole mass spectrometer (Hewlett Packard, Palo Alto, California). CHC extracts of wasps studied at the field site Tenneville in 2017 were analyzed with a gas chromatograph-flame ionization detector (Shimadzu GC-2010 system, Shimadzu, Wemmel, Belgium) equipped with a SLB-5 MS non-polar capillary column (5%-phenyl)-methylpolysiloxane stationary phase; 30 m × 0.25 mm ID; film thickness: 0.25 µm). We consistently applied the following temperature profile when analyzing CHC extracts of wasps: start temperature 60 °C, temperature increase of 20 °C per minute up to 150 °C, followed by a temperature increase of 5 °C per minute up to 300 °C, which was maintained for 10 min. Honey bee CHC extracts were analyzed with the Agilent GC-MS specified above and applying the following temperature profile: start temperature 60 °C, followed by a temperature increase of 5 °C per minute up to 300 °C, which was maintained for 10 min.

*Genomic DNA NGS libraries*

We generated three genomic DNA NGS libraries from a single pool of DNA extracted from meso- and metasomas of ten males (collected May 15 and 16, 2013) and 31 females (collected May 15 and 16, 2013) of *Odynerus spinipes* collected near Würzburg, Germany (49.77977, 9.97065) (voucher samples of the female samples are available in the BioBank of Oliver Niehuis at the University of Freiburg: ON_0890–0899, ON_0923, ON_0924, ON_2277–ON_2295).

*Raw read processing*

All raw reads of genomic DNA were processed with trimmomatic[46] version 0.33, selecting paired-end mode (if applicable), the Illumina TruSeq3 adapter sequences provided with trimmomatic, and the parameters: LEADING:3, TRAILING:3, SLIDINGWINDOW: 4:25, MINLEN:50. The RNA-seq raw reads were processed with the same software.

*Masking of transposable elements and annotation of protein-coding genes*

We used RepeatModeler version 1.0.8 (http://www.repeatmasker.org) to compile a *de novo* repeat library. The pipeline identifies transposon (TE) nucleotide sequences by comparing the *de novo* repeat library to entries in the NCBI nr database (downloaded 03/17/2016 from https://ftp.ncbi.nlm.nih.gov/blast/db/) and removing nucleotide sequences that are not associated with transposons. We applied the same filter parameters as used by Petersen et al.[71]. The pipeline further combines the filtered repeat library with the Metazoa-specific section of RepBase[72] version 20140131 to create the final repeat library. RepeatMasker[73] version 4.0.5 was then used to annotate TEs in the *O. spinipes* genome assembly and to generate a soft-masked version of the draft genome. Protein-coding genes were annotated with the BRAKER[48] *ab initio* gene prediction pipeline version 2.1. To this end, we first mapped the available trimmed RNA-seq raw reads onto the soft-masked version of the *O. spinipes* draft genome assembly using HISAT2[74,75] version 2.1.0 and converted the output into a sorted BAM file using SAMtools[76] version 1.7 and BAMtools[77] version 2.3.0. The resulting BAM file was, together with the soft-masked draft genome assembly, provided as input to the BRAKER2 pipeline (internally using GeneMark[78] version 4.33; Augustus[79] version 3.3; NCBI BLAST+[58,59] version 2.6.0; SAMtools[76] version 1.7; and BAMtools[77] version 2.5.1) to predict protein-coding genes. BRAKER2 parameters (apart from path specifications) were set as follows: --UTR=off --gff3 --softmasking. UTR annotation was disabled, as it was at the time of execution according to the documentation of Augustus 3.3 not suitable for annotation of non-model insect genomes.

*Gene tree taxon sampling*

We searched the genomes of the following 36 Euarthropoda for genes/proteins of gene families of interest: *Acromyrmex echinatior* (OGS version 3.8[80]), *Aedes aegypti* (gene set version AaegL3.3[81]), *Amyelois transitella* (gene set version 1; NCBI GCA_001186105.1), *Apis mellifera* (OGS version 3.2[82]), *Bombus terrestris* (gene set version 1.0[83]), *Bombyx mori* (gene set version 1.0[84]), *Calopteryx splendens* (OGS version 1.0[85]), *Camponotus floridanus* (gene set version 3.3[86]), *Catajapyx aquilonaris* (gene set Caqu_2.0, NCBI GCA_000934665.2[87]), *Centruroides sculpturatus* (OGS version 2.0[88]), *Cimex lectularius* (OGS version 1.2[89]), *Daphnia pulex* (OGS version 1.0[90]), *Dendroctonus ponderosae* (OGS version 1.0[91]), *Drosophila melanogaster* (OGS version r6.03[92]), *Drosophila simulans* (OGS version r1.4[93]), *Eriocheir sinensis* (gene set version 1.0; NCBI GCA_013436485.1[94]), *Eurytemora affinis* (gene set version AFF_2.0; NCBI GCA_000591075.2[95]), *Harpegnathos saltator* (OGS version 3.3[86]), *Hyalella azteca* (gene set version Hazt_2.0[96]), *Ixodes scapularis* (gene set version JCVI_ISG_i3_1.0[97]), *Latrodectus hesperus* (gene set version Lhes_2.0; NCBI GCA_000697925.2), *Lepeophtheirus salmonis* (gene set version LSal-F-Atl-CAN-001; NCBI 001005205.1), *Limulus polyphemus* (gene set version 2.1.2; NCBI GCF_000517525.1[98]), *Macrotermes natalensis* (gene set version 1.2[99]), *Manduca sexta* (OGS version 2[100]), *Nasonia vitripennis* (OGS version 2.26[101]), *Parasteatoda tepidariorum* (gene set version aug3.1[88]), *Pediculus humanus* (gene set version U2.1[102]), *Rhodnius prolixus* (gene set version. C1.2[103]), *Sarcoptes scabiei* (gene set version SscaA1[104]), *Stegodyphus mimosarum* (gene set version 1; NCBI GCA_000611955.2[105]), *Strigamia maritima* (gene set version Smar1[106]), *Tetranychus urticae* (gene set version 1.0[107]), *Tribolium castaneum* (OGS version Tcas2.0[108]), *Tigriopus californicus* (gene set version Tcal_SD_v2.1; NCBI GCA_007210705.1), *Zootermopsis nevadensis* (OGS version 2.2[109]).

*Gene tree inference*

The amino acid sequences of a given gene family were aligned with MAFFT[56] version 7.123 and applying the run parameters "--maxiterate 1000 --globalpair –reorder". The resulting multiple amino acid sequence alignments were analyzed with IQ-TREE[57] version

1.6 to infer a gene tree of each gene family under the maximum likelihood optimality criterion. We used the corrected Akaike information criterion (AICc) to select the best-fitting amino acid substitution model for each gene family separately (IQ-TREE option: "-madd LG4M, LG4X"). Branch support was inferred by using 1) ultrafast bootstrapping (UFBoot2) and specifying the run parameters "-bb" and "-bnni" and 2) Shimodaira-Hasegawa approximate likelihood ratio tests ([SH]-aLRT; IQ-TREE option: -alrt 10000)[110]. Branch support was considered reliable if SH-aLRT was larger or equal 80 % and UFboot was larger or equal 95 %. We used FigTree version 1.4.3 (http://tree.bio.ed.ac.uk/software/figtree/) to midway-root the inferred gene trees. Because FigTree is known to map in some situations branch support values to wrong branches when re-rooting a tree[111], we visually verified all bootstrap values with the Interactive Tree of Life (ITOL)[112] version 3 online tool. The gene trees were further processed and annotated, using information from FlyAtlas2[113], with Inkscape 1.1 (http://www.inkscape.org/).

*Modifications applied for whole mount in situ hybridization*

Whole-mount RNA *in situ* hybridization was done following the protocol for staining genes in brains and ovaries of adults given by [63] with the following modifications: step 14, probes were diluted in 300 $\mu$L (not in 50 $\mu$L); step 17, tissues were kept in wash buffer at 57 °C for 1 h (not overnight); step 20, we used the anti-Fluorescein-AP Fab fragment [150 Units] (Roche, Mannheim, Germany) or the anti-Digoxigenin-AP Fab fragment [150 Units] (Roche, Mannheim, Germany) antibodies depending on the riboprobes and on the RNA labelling mix used to synthesize it. Stained samples were rehydrated into PBS, dissected and mounted before imaging.

*Gene selection for knockdown experiments*

Genes selected for knockdown experiments were found expressed in oenocytes (except GB51238, see below), showed a high absolute *log*$_2$-fold change value (*Supplementary*

*Tables 7*, *8*, and *9*), and were chosen to represent a variety of gene families (we tried to investigate at least one gene per gene families). We decided to investigate desaturase GB51238, even though it was found to be expressed in trophocytes, since it was one of the three co-orthologs of the gene candidate g14712. While the other two genes are expressed in oenocytes, we intentionally decided to select GB51238 to assess a possible involvement also of trophocytes in CHC biosynthesis.

*Double-stranded RNA synthesis*

We synthesized dsRNA from the same 387–774-bp-long DNA fragments that we had cloned to facilitate anti-sense riboprobe synthesis. Specifically, template DNA for dsRNA synthesis was generated by PCR-amplifying with the Phusion High Fidelity kit (New England Biolabs, Ipswich, MA, USA) the plasmid inserts using an oligonucleotide primer listed for the specific gene in *Supplementary Table 10* at one end and a T7 oligonucleotide primer at the other end of the plasmid insert. By reversing the order of the oligonucleotides, we obtained dsRNA template DNA with the T7 promotor sequence on different sides of the amplicon.

**Reference supplementary Methods**


71. Petersen, M., Armisén, D., Gibbs, R. A., Hering, L., Khila, A., Mayer, G., Richards, S., Niehuis, O., Misof, B. Diversity and evolution of the transposable element repertoire in arthropods with particular reference to insects. *BMC Evol. Biol.* **19**, 1–15 (2019).

72. Jurka, J., Kapitonov, V. V., Pavlicek, A., Klonowski, P., Kohany, O., Walichiewicz, J. Repbase Update, a database of eukaryotic repetitive elements. *Cytogenet. Genome Res.* **110**, 462–467 (2005). Accessed 1 Sept 2016.

73. Bailly-Bechet M., Haudry A., Lerat E. One code to find them all: a Perl tool to conveniently parse RepeatMasker output files. *Mob. DNA* **5**, 13 (2014).

74. Kim, D., Langmead, B. & Salzberg, S. L. HISAT: a fast spliced aligner with low memory requirements. *Nat. Methods* **12**, 357–360 (2015).

75. Kim, D., Paggi, J.M., Park, C. et al. Graph-based genome alignment and genotyping with HISAT2 and HISAT-genotype. *Nat. Biotechnol.* **37**, 907–915 (2019).

76. Li, H., Handsaker, B., Wysoker, A., Fennell, T., Ruan, J., Homer, N., Marth, G., Abecasis, G. & Durbin, R. The sequence alignment/map format and SAMtools. *Bioinformatics* **25**, 2078–2079 (2009).

77. Barnett, D. W., Garrison, E. K., Quinlan, A. R., Strömberg, M. P. & Marth, G. T. BamTools: a C++ API and toolkit for analyzing and managing BAM files. *Bioinformatics* **27**, 1691–1692 (2011).

78. Besemer, J. & Borodovsky, M. GeneMark: web software for gene finding in prokaryotes, eukaryotes and viruses. *Nucleic acids Res.* **33**, W451–W454 (2005).

79. Stanke, M., Steinkamp, R., Waack, S. & Morgenstern, B. AUGUSTUS: a web server for gene finding in eukaryotes. *Nucleic acids Res.* **32**, W309–W312 (2004).

80. Nygaard, S., Zhang, G., Schiøtt, M., Li, C., Wurm, Y., Hu, H., et al. The genome of the leaf-cutting ant *Acromyrmex echinatior* suggests key adaptations to advanced social life and fungus farming. *Genome Res.* **21**, 1339–1348 (2011).

81. Nene, V., Wortman, J. R., Lawson, D., Haas, B., Kodira, C., Tu, Z. J., et al. Genome sequence of *Aedes aegypti*, a major arbovirus vector. *Science* **316**, 1718–1723 (2007).

82. Elsik, C. G., Worley, K. C., Bennett, A. K., Beye, M., Camara, F., Childers, C. P., de Graaf, D. C., Debyser, G., Deng, J., Devreese, B., et al. Finding the missing honey bee genes: lessons learned from a genome upgrade. *BMC Genomics* **15**, 1–29 (2014).

83. Sadd, B. M., Barribeau, S. M., Bloch, G., De Graaf, D. C., Dearden, P., Elsik, C. G., et al. The genomes of two key bumblebee species with primitive eusocial organization. *Genome Biol.* **16**, 1–32 (2015).

84. International Silkworm Genome Consortium. The genome of a lepidopteran model insect, the silkworm *Bombyx mori*. *Insect Biochem. Mol. Biol.* **38**, 1036–1045 (2008).

85. Ioannidis, P., Simao, F. A., Waterhouse, R. M., Manni, M., Seppey, M., Robertson, H. M., Misof, B., Niehuis, O., Zdobnov, E. M. Genomic features of the damselfly *Calopteryx splendens* representing a sister clade to most insect orders. *Genome Biol. Evol.* **9**, 415–430 (2017).

86. Bonasio, R., Zhang, G., Ye, C., Mutti, N. S., Fang, X., Qin, N., Donahue, G., Yang, P., Li, Q., Li, C., Zhang, P., Huang, Z., Berger, S. L., Reinberg, D., Wang, J., Liebig, J. Genomic comparison of the ants *Camponotus floridanus* and *Harpegnathos saltator*. *Science* **329**,1068–1071 (2010).

87. Thomas, G. W. C., Dohmen, E., Hughes, D. S. T., Murali, S. C., Poelchau, M., Glastad, K., Anstead, C. A., Ayoub, N. A., Batterham, P., Bellair, M. et al. Gene content evolution in the arthropods. *Genome Biol.* **21**, 15 (2020).

88. Schwager, E. E., Sharma, P. P., Clarke, T., Leite, D. J., Wierschin, T., Pechmann, M., et al. The house spider genome reveals an ancient whole-genome duplication during arachnid evolution. *BMC Biol.* **15**, 1–27 (2017).

89. Rosenfeld, J. A., Reeves, D., Brugler, M. R., Narechania, A., Simon, S., Durrett, R., et al. Genome assembly and geospatial phylogenomics of the bed bug *Cimex lectularius*. *Nat. Commun.* **7**, 1–10 (2016).

90. Colbourne, J. K., Pfrender, M. E, Gilbert, D., Thomas, W. K., Tucker, A., Oakley, T. H., Tokishita, S., Aerts, A., Arnold, G. J., Basu, M. K., et al. The ecoresponsive genome of *Daphnia pulex*. *Science* **331**, 555–561 (2011).

91. Keeling, C. I., Yuen, M. M., Liao, N. Y., Docking, T. R., Chan, S. K., Taylor, G. A., et al. Draft genome of the mountain pine beetle, *Dendroctonus ponderosae* Hopkins, a major forest pest. *Genome Biol.* **14**, 1–20 (2013).

92. Adams, M. D., Celniker, S. E., Holt, R. A., Evans, C. A., Gocayne, J. D., Amanatides, P. G., Scherer, S. E., Li, P.W. et al. The genome sequence of *Drosophila melanogaster*. *Science* **287**, 2185–2195 (2000).

93. Hu, T. T., Eisen, M. B., Thornton, K. R., Andolfatto, P. A second-generation assembly of the *Drosophila simulans* genome provides new insights into patterns of lineage-specific divergence. *Genome Res*. **23**, 89–98 (2013).

94. Cui, Z., Liu, Y., Yuan, J., Zhang, X., Ventura, T., Ma, K. Y., Sun, S., Song, C., Zhan, D., Yang, Y. et al. The Chinese mitten crab genome provides insights into adaptive plasticity and developmental regulation. *Nat. Commun*. **12**, 2395 (2021).

95. Choi, B. S., Kim, D. H., Kim, M. S., Park, J. C., Lee, Y. H., Kim, H. J., Jeong, C. B., Hagiwara, A., Souissi, S., Lee, J. S. The genome of the European estuarine calanoid copepod *Eurytemora affinis*: Potential use in molecular ecotoxicology. *Mar. Pollut. Bull.* **166**, 112190 (2021).

96. Poynton, H. C., Hasenbein, S., Benoit, J. B., Sepulveda, M. S., Poelchau, M. F., Hughes, D. S., et al. The toxicogenome of *Hyalella azteca*: a model for sediment ecotoxicology and evolutionary toxicology. *Environ. Sci. Technol.* **52**, 6009–6022 (2018).

97. Gulia-Nuss, M., Nuss, A. B., Meyer, J. M., Sonenshine, D. E., Roe, R. M., Waterhouse, R. M., et al. Genomic insights into the *Ixodes scapularis* tick vector of Lyme disease. *Nat. Commun.* **7**, 1–13 (2016).

98. Simpson, S. D., Ramsdell, J. S., Watson III, W. H. & Chabot, C. C. (2017). The draft genome and transcriptome of the Atlantic horseshoe crab, *Limulus polyphemus*. *Int. J. Genomics* **2017**, 7636513 (2017).

99. Poulsen, M., Hu, H., Li, C., Chen, Z., Xu, L., Otani, S., et al. Complementary symbiont contributions to plant decomposition in a fungus-farming termite. *Proc. Natl. Acd. Sci. USA* **111**, 14500–14505 (2014).

100. Kanost, M. R., Arrese, E. L., Cao, X., Chen, Y. R., Chellapilla, S., Goldsmith, M. R., et al. Multifaceted biological insights from a draft genome sequence of the tobacco hornworm moth, *Manduca sexta*. *Insect Biochem. Mol. Biol.* **76**, 118–147 (2016).

101. Werren, J. H., Richards, S., Desjardins, C. A., Niehuis, O., Gadau, J., Colbourne, J. K. et al. Functional and evolutionary insights from the genomes of three parasitoid Nasonia species. *Science* **327**, 343–348 (2010).

102. Kirkness, E. F., Haas, B. J., Sun, W., Braig, H. R., Perotti, M. A., Clark, J. M., et al. Genome sequences of the human body louse and its primary endosymbiont provide insights into the permanent parasitic lifestyle. *Proc. Natl. Acad. Sci. USA* **107**, 12168–12173 (2010).

103. Mesquita, R. D., Vionette-Amaral, R. J., Lowenberger, C., Rivera-Pomar, R., Monteiro, F. A., Minx, P., et al. Genome of *Rhodnius prolixus*, an insect vector of Chagas disease, reveals unique adaptations to hematophagy and parasite infection. *Proc. Natl. Acad. Sci. USA* **112**, 14936–14941 (2015).

104. Rider, S. D., Morgan, M. S. & Arlian, L. G. Draft genome of the scabies mite. *Parasit. Vectors* **8**, 1–14 (2015).

105. Sanggaard, K. W., Bechsgaard, J. S., Fang, X., Duan, J., Dyrlund, T. F., Gupta, V., et al. Spider genomes provide insight into composition and evolution of venom and silk. *Nat. Commun.* **5**, 1–12 (2014).

106. Chipman, A. D., Ferrier, D. E., Brena, C., Qu, J., Hughes, D. S., Schröder, R., Torres-Oliva, M., Znassi, N., Jiang, H., Almeida, F. C., Alonso, C. R., Apostolou, Z. et al. The first myriapod genome sequence reveals conservative arthropod gene content and genome organisation in the centipede *Strigamia maritima*. *PLoS Biol.* **12**, e1002005 (2014).

107. Grbić, M., Van Leeuwen, T., Clark, R. M., Rombauts, S., Rouzé, P., Grbić, V., et al. The genome of *Tetranychus urticae* reveals herbivorous pest adaptations. *Nature* **479**, 487–492 (2011).

108. Tribolium Genome Sequencing Consortium. The genome of the model beetle and pest *Tribolium castaneum*. *Nature* **452**, 949–955 (2008).

109. Terrapon, N., Li, C., Robertson, H. M., Ji, L., Meng, X., Booth, W., Chen, Z., Childers, C. P., Glastad, K. M., Gokhale, K., Gowin, J., Gronenberg, W. et al. Molecular traces of alternative social organization in a termite genome. *Nat. Commun*. **5**, 3636 (2014).

110. Guindon, S., Dufayard, J. F., Lefort, V., Anisimova, M., Hordijk, W. & Gascuel, O. New algorithms and methods to estimate maximum-likelihood phylogenies: assessing the performance of PhyML 3.0. *Syst. Biol.* **59**, 307–321 (2010).

111. Czech, L., Huerta-Cepas, J. & Stamatakis, A. A critical review on the use of support values in tree viewers and bioinformatics toolkits. *Mol. Biol. Evol.* **34**, 1535–1542 (2017).

112. Letunic, I. & Bork, P. Interactive tree of life (iTOL) v3: an online tool for the display and annotation of phylogenetic and other trees. *Nucleic Acids Res.* **44**, W242–W245 (2016).

113. Leader, D. P., Krause, S. A., Pandit, A., Davies, S. A. & Dow, J. A. T. FlyAtlas 2: a new version of the Drosophila melanogaster expression atlas with RNA-Seq, miRNA-Seq and sex-specific data. *Nucleic Acids Res.* **46**, D809–D815 (2018).

### Supplementary Tables

***Supplementary Table 1***. Sample dates of *Odynerus spinipes* females, kept in the laboratory and whose cuticular hydrocarbon profile (chemotype) we studied multiple times over the course of their adult life.

| Sample ID | Chemo-type | Eclosion date | 1st sampling date | Age at 1st sampling date | 2nd sampling date | Age at 2nd sampling date | 3rd sampling date | Age at 3rd sampling date |
|---|---|---|---|---|---|---|---|---|
| ON_8469[a] | 1 | April 25, 2016 | April 26, 2016 | 1 | May 4, 2016 | 9 | May 11, 2016 | 16 |
| ON_8471[b] | 2 | April 25, 2016 | April 26, 2016 | 1 | May 2, 2016 | 7 | May 9, 2016 | 14 |
| ON_8473 | 2 | April 25, 2016 | April 27, 2016 | 2 | May 2, 2016 | 7 | May 10, 2016 | 15 |
| ON_8475[c] | 1 | April 25, 2016 | April 26, 2016 | 1 | May 2, 2016 | 7 | NA | NA |
| ON_8477[d] | 2 | April 25, 2016 | April 26, 2016 | 1 | May 4, 2016 | 9 | NA | NA |
| ON_8479 | 1 | April 25, 2016 | April 27, 2016 | 2 | May 2, 2016 | 7 | May 10, 2016 | 15 |
| ON_8481 | 1 | April 25, 2016 | April 27, 2016 | 2 | May 2, 2016 | 7 | May 10, 2016 | 15 |
| ON_8483 | 1 | April 25, 2016 | April 27, 2016 | 2 | May 2, 2016 | 7 | May 10, 2016 | 15 |
| ON_8485 | 2 | April 25, 2016 | April 27, 2016 | 2 | May 3, 2016 | 8 | May 9, 2016 | 14 |
| ON_8487 | 1 | April 25, 2016 | April 27, 2016 | 2 | May 2, 2016 | 7 | NA | NA |
| ON_8489 | 2 | April 25, 2016 | April 27, 2016 | 2 | May 3, 2016 | 8 | May 10, 2016 | 15 |
| ON_8491 | 2 | April 26, 2016 | April 27, 2016 | 1 | May 3, 2016 | 7 | May 9, 2016 | 13 |
| ON_8493 | 2 | April 26, 2016 | April 28, 2016 | 2 | May 3, 2016 | 7 | May 10, 2016 | 14 |
| ON_8495 | 2 | April 26, 2016 | April 28, 2016 | 2 | May 2, 2016 | 6 | NA | NA |
| ON_8497 | 2 | April 26, 2016 | April 28, 2016 | 2 | May 2, 2016 | 6 | NA | NA |
| ON_8499 | 1 | April 26, 2016 | April 28, 2016 | 2 | May 3, 2016 | 7 | May 9, 2016 | 13 |
| ON_8501 | 2 | April 26, 2016 | April 28, 2016 | 2 | May 3, 2016 | 7 | May 9, 2016 | 13 |
| ON_8503 | 2 | April 26, 2016 | April 28, 2016 | 2 | May 3, 2016 | 7 | May 10, 2016 | 14 |
| ON_8505 | 2 | April 26, 2016 | April 28, 2016 | 2 | May 3, 2016 | 7 | May 9, 2016 | 13 |
| ON_8507 | 2 | April 27, 2016 | April 29, 2016 | 2 | May 4, 2016 | 7 | May 10, 2016 | 13 |
| ON_8509 | 1 | April 27, 2016 | April 29, 2016 | 2 | May 2, 2016 | 5 | NA | NA |
| ON_8511 | 2 | April 27, 2016 | April 29, 2016 | 2 | May 4, 2016 | 7 | May 10, 2016 | 13 |
| ON_8513 | 1 | April 27, 2016 | April 29, 2016 | 2 | May 4, 2016 | 7 | May 11, 2016 | 14 |
| ON_8515 | 2 | April 27, 2016 | April 29, 2016 | 2 | May 4, 2016 | 7 | May 11, 2016 | 14 |
| ON_8517 | 2 | April 27, 2016 | April 29, 2016 | 2 | May 4, 2016 | 7 | May 11, 2016 | 14 |
| ON_8519 | 1 | April 27, 2016 | April 28, 2016 | 1 | May 4, 2016 | 7 | May 11, 2016 | 14 |
| ON_8521 | 1 | April 28, 2016 | April 29, 2016 | 1 | NA | NA | NA | NA |
| ON_8523 | 2 | April 27, 2016 | April 29, 2016 | 2 | May 4, 2016 | 7 | May 11, 2016 | 14 |
| ON_8525 | 2 | April 28, 2016 | April 29, 2016 | 1 | NA | NA | NA | NA |
| ON_8527 | 1 | April 28, 2016 | April 29, 2016 | 1 | NA | NA | NA | NA |

[a] Extracurricular sampling at April 29, 2016 (wasp's age: 4 days)
[b] Extracurricular sampling at April 27, 2016 (wasp's age: 3 days)
[c] Extracurricular sampling at April 27, 2016 (wasp's age: 2 days)
[c] Extracurricular sampling at April 29, 2016 (wasp's age: 4 days)

***Supplementary Table 2***. Sample dates of *Odynerus spinipes* females whose cuticular hydrocarbon profile (chemotype) we studied multiple times over the course of their adult life at two field sites (Büchelberg [B; 49.027985, 8.164801] and Tenneville [T; 50.100671, 5.530061]).

| Sample ID | Chemo-type | 1st sampling date | Idle time (days) | 2nd sampling date | Idle time (days) | 3rd sampling date | Idle time (days) | 4th sampling date | Field site |
|---|---|---|---|---|---|---|---|---|---|
| ON_8651 | 1 | June 7, 2016 | 8 | June 15, 2016 | NA | NA | NA | NA | B |
| ON_8653 | 1 | June 7, 2016 | 13 | June 20, 2016 | NA | NA | NA | NA | B |
| ON_8663 | 1 | June 15, 2016 | 5 | June 20, 2016 | NA | NA | NA | NA | B |
| ON_8671 | 2 | May 27, 2017 | 4 | Mai 31, 2017 | 8 | June 8, 2017 | NA | NA | B |
| ON_8673 | 1 | May 27, 2017 | 6 | June 2, 2017 | 6 | June 8, 2017 | 11 | June 19, 2017 | B |
| ON_8675 | 1 | May 31, 2017 | 8 | June 8, 2017 | 11 | June 19, 2017 | NA | NA | B |
| ON_8677 | 2 | May 31, 2017 | 8 | June 8, 2017 | NA | NA | NA | NA | B |
| ON_8693 | 1 | June 10, 2017 | 3 | June 13, 2017 | 2 | June 15, 2017 | NA | NA | B |
| ON_8699 | 2 | June 10, 2017 | 3 | June 13, 2017 | NA | NA | NA | NA | B |
| ON_8709 | 2 | June 13, 2017 | 6 | June 19, 2017 | 3 | June 21, 2017 | NA | NA | B |
| ON_8721 | 2 | June 21, 2017 | 2 | June 23, 2017 | NA | NA | NA | NA | B |
| ON_8723 | 2 | June 21, 2017 | 2 | June 23, 2017 | NA | NA | NA | NA | B |
| ON_8725 | 1 | July 3, 2017 | 2 | July 5, 2017 | NA | NA | NA | NA | T |
| ON_8729 | 2 | July 3, 2017 | 2 | July 5, 2017 | NA | NA | NA | NA | T |
| ON_8733 | 2 | July 3, 2017 | 3 | July 6, 2017 | NA | NA | NA | NA | T |
| ON_8735 | 2 | July 3, 2017 | 3 | July 6, 2017 | NA | NA | NA | NA | T |
| ON_8737 | 2 | July 4, 2017 | 2 | July 6, 2017 | NA | NA | NA | NA | T |
| ON_8745 | 1 | July 4, 2017 | 2 | July 6, 2017 | NA | NA | NA | NA | T |

***Supplementary Table 3***. Information to *Odynerus spinipes* transcriptomes we sequenced and used to facilitate annotation of the *Odynerus spinipes* draft genome.

| Sample ID | Tissue ID | Sex | Chemo-type | Tissue | Sampling location | Collection date |
|---|---|---|---|---|---|---|
| ON_6860 | Odsp_RNA_05 | female | 1 | whole body | Büchelberg | May 19, 2014 |
| ON_6859 | Odsp_RNA_09 | female | 2 | whole body | Büchelberg | May 19, 2014 |
| ON_6846 | Odsp_RNA_08 | male | NA | whole body | Büchelberg | May 18, 2014 |

***Supplementary Table 4***. Adult *Odynerus spinipes* females whose metasoma transcriptomes we sequenced and which we collected near Bad Muskau (51.548200, 14.714900), Büchelberg (49.027985, 8.164801), and Eichenzell (50.495927, 9.704589).

| **Sample ID** | **Chemo-type** | **Collection site** | **RNA fixation date** | **Age class** | **Batch no.** |
|---|---|---|---|---|---|
| ON_7724 | 1 | Büchelberg | January 27, 2015 | 12–38 h | 1 |
| ON_7728 | 1 | Büchelberg | January 29, 2015 | 12–38 h | 1 |
| ON_7747 | 1 | Büchelberg | February 12, 2015 | 12–38 h | 1 |
| ON_7723 | 2 | Büchelberg | January 27, 2015 | 12–38 h | 1 |
| ON_7730 | 2 | Büchelberg | January 29, 2015 | 12–38 h | 1 |
| ON_7739 | 2 | Büchelberg | January 31, 2015 | 12–38 h | 1 |
| ON_8683 | 1 | Büchelberg | June 3, 2017 | 48–62 h | 2 |
| ON_8685 | 1 | Büchelberg | June 3, 2017 | 48–62 h | 2 |
| ON_12769 | 1 | Eichenzell | June 12, 2016 | 48–62 h | 2 |
| ON_8687 | 2 | Büchelberg | June 6, 2017 | 48–62 h | 2 |
| ON_12771 | 2 | Eichenzell | June 14, 2016 | 48–62 h | 2 |
| ON_12775 | 2 | Bad Muskau | May 17, 2017 | 48–62 h | 2 |

***Supplementary Table 5***. *Odynerus spinipes* genes significantly differentially expressed in batch 1-females with different chemotype. The table shows the gene IDs, the *log*$_2$-fold change values (values < 0 indicate higher expression of genes in females with chemotype 2 than in females with chemotype 1), the *p*-value obtained with DESeq2, the adjusted *p*-value from applying the false discovery rate (FDR), and the predicted function of the genes based on homology.

*Provided as an excel table "Supplementary_table_5.xlsx"*

***Supplementary Table 6***. *Odynerus spinipes* genes significantly differentially expressed in batch 2-females with different chemotype. The table shows the gene IDs, the *log*$_2$-fold change values (values < 0 indicate higher expression of genes in females with chemotype 2 than in females with chemotype 1), the *p*-value obtained with DESeq2, the adjusted *p*-value from applying the false discovery rate (FDR), and the predicted function of the genes based on homology.

*Provided as an excel table "Supplementary_table_6.xlsx"*

***Supplementary Table 7***. *Odynerus spinipes* genes significantly differentially expressed in females with different chemotype in two batches of samples differing in age and sampling location from each other (***Supplementary Table 4***). The table shows the results from first globally assessing gene expression differences in batch 1 and subsequently assessing whether those genes that are considered differentially expressed based on false discovery rate (FDR) are also differentially expressed (*p*-value ≤ 0.05) in batch 2 after applying Holm-Bonferroni correction for multiple testing (number of genes tested in batch 2) on the *p*-values provided by DESeq2. $Log_2$-fold change values > 0 indicate higher expression of genes in females with chemotype 1 than in females with chemotype 2. $Log_2$-fold change values < 0 indicate higher expression of genes in females with chemotype 2 than in females with chemotype 1. Genes in bold were significantly differently expressed in both batches.

| Gene ID | Batch 1 | | | Batch 2 | | | Predicted function |
|---|---|---|---|---|---|---|---|
| | $Log_2$-fold change | *p* | *p* (adj.)[1] | $Log_2$-fold change | *p* | *p* (adj.)[2] | |
| **g283** | -0.96 | 0.000 | 0.003 | -1.73 | 0.000 | **0.000** | **rhythmically expressed gene 5 protein** |
| g1571 | -1.27 | 0.000 | 0.000 | -1.09 | 0.031 | 1.000 | fatty acid reductase |
| **g2290** | -0.84 | 0.000 | 0.029 | -1.79 | 0.000 | **0.019** | **fatty acid amine hydrolase** |
| g3041 | 1.13 | 0.000 | 0.000 | -0.89 | 0.032 | 1.000 | serine-rich adhesin for platelets |
| **g3059** | -1.55 | 0.000 | 0.000 | -1.71 | 0.000 | **0.013** | **parathyroid hormone/related receptor** |
| g3158 | -1.79 | 0.000 | 0.000 | -1.00 | 0.024 | 1.000 | fatty acid synthase |
| g4709 | -0.91 | 0.000 | 0.043 | -1.14 | 0.021 | 1.000 | alpha-tocopherol transfer protein-like |
| g6414 | -1.04 | 0.000 | 0.003 | -1.19 | 0.018 | 0.968 | alanine--glyoxylate aminotransferase 2 homolog 1, mitochondrial |
| **g7616** | -1.19 | 0.000 | 0.000 | -1.46 | 0.000 | **0.008** | **fatty acid elongase** |
| g12825 | 0.75 | 0.000 | 0.049 | 1.01 | 0.002 | 0.090 | adenylate kinase isoenzyme 5 |
| g14547 | -2.54 | 0.000 | 0.000 | -1.09 | 0.018 | 0.968 | retrovirus-related pol polyprotein from transposon tnt 1-94 |
| **g14712** | -0.99 | 0.000 | 0.008 | -3.85 | 0.000 | **0.000** | **fatty acid desaturase** |

[1] FDR

[2] Holm-Bonferroni correction for 60 tests

***Supplementary Table 8***. *Odynerus spinipes* genes significantly differentially expressed in females with different chemotype in two batches of samples differing in age and sampling location from each other (***Supplementary Table 4***). The table shows the results from first globally assessing gene expression differences in batch 2 and subsequently assessing whether those genes that are considered differentially expressed based on false discovery rate (FDR) are also differentially expressed (*p*-value ≤ 0.05) in batch 1 when applying Holm-Bonferroni correction for multiple testing (number of genes tested in batch 1) on the *p*-values provided by DESeq2. *Log₂*-fold change values < 0 indicate higher expression of genes in females with chemotype 2 than in females with chemotype 1. Genes in bold were significantly differently expressed in both batches.

| Gene ID | Batch 2 | | | Batch 1 | | | Predicted function |
|---|---|---|---|---|---|---|---|
| | ***Log$_2$*-fold change** | ***p*** | ***p* (adj.)[1]** | ***Log$_2$*-fold change** | ***p*** | ***p* (adj.)[2]** | |
| **g283** | -1.73 | 0.000 | 0.004 | -0.96 | 0.000 | **0.000** | **rhythmically expressed gene 5 protein** |
| g531 | -1.70 | 0.000 | 0.013 | -0.49 | 0.025 | 0.685 | progestin and adipoQ receptor family member 3 isoform X1 |
| g7599 | 1.92 | 0.000 | 0.032 | -0.51 | 0.048 | 1.000 | NA |
| **g7616** | -1.46 | 0.000 | 0.049 | -1.19 | 0.000 | **0.000** | **fatty acid elongase** |
| g14676 | -1.75 | 0.000 | 0.008 | -0.52 | 0.024 | 0.665 | 1-acyl-sn-glycerol-3-phosphate acyltransferase alpha |
| **g14712** | -3.85 | 0.000 | 0.000 | -0.99 | 0.000 | **0.000** | **fatty acid desaturase** |

[1] FDR
[2] Holm-Bonferroni correction for 31 tests

***Supplementary Table 9***. *Odynerus spinipes* genes encoding fatty acid synthases, fatty acid elongases, fatty acid desaturases, and fatty acid reductases and found significantly differentially expressed in females belonging to different age classes (***Supplementary Table 4***). Shown are the results from analyzing the twelve transcriptomes with the two software packages DESeq2 and EdgeR. *Log$_2$*-fold change values > 0 indicate higher expression of genes in 48–62-h-old females than in 12–38-h-old females. *Log$_2$*-fold change values < 0 indicates higher expression of genes in 12–38-h-old females than in 48–62-h-old females. FDR: false discovery rate.

| Gene ID | DESeq2 | | | EdgeR | | | Gene family |
|---|---|---|---|---|---|---|---|
| | ***log$_2$*-fold change** | ***p*** | **FDR** | ***log$_2$*-fold change** | ***p*** | **FDR** | |
| g1571 | 3.170 | 0.001 | **0.001** | 4.691 | 0.000 | **0.000** | fatty acid reductase |
| g2413 | -0.767 | 0.003 | **0.003** | -0.813 | 0.002 | **0.010** | fatty acid reductase |
| g3158 | 1.972 | 0.017 | **0.017** | 4.361 | 0.003 | **0.012** | fatty acid synthase |
| g6537 | 1.482 | 0.014 | **0.014** | 1.739 | 0.006 | **0.022** | fatty acid elongase |
| g7609 | NA | NA | NA | 2.711 | 0.001 | **0.004** | fatty acid elongase |
| g7610 | NA | NA | NA | 7.095 | 0.000 | **0.000** | fatty acid elongase |
| g7616 | 1.454 | 0.004 | **0.004** | 1.602 | 0.002 | **0.009** | fatty acid elongase |
| g7620 | 1.449 | 0.036 | **0.036** | NA | NA | NA | fatty acid elongase |
| g7621 | NA | NA | NA | 3.535 | 0.001 | **0.005** | fatty acid elongase |
| g7842 | 1.208 | 0.004 | **0.004** | 1.288 | 0.001 | **0.005** | fatty acid reductase |
| g14708 | -0.691 | 0.013 | **0.013** | -0.730 | 0.004 | **0.016** | fatty acid desaturase |

***Supplementary Table 10***. Number of samples from which cuticular hydrocarbon and gene expression data were collected in RNAi experiments. The time between dsRNA injection and data collection is specified in the column with the header **Δ***t.*

| Honey bee gene ID | *O. spinipes* gene ID | Predicted function | Δ*t* [days] | Treatment group (RT-qPCR) sample size | Control group (RT-qPCR) sample size |
|---|---|---|---|---|---|
| GB40659 | g14708 | fatty acid desaturase | 2 | 7 (NA) | 6 (NA) |
| GB42218 | g14712_18 | fatty acid desaturase | 4 | 12 (12) | 13 (11) |
| GB48195 | g14710 | fatty acid desaturase | 3 | 11 (8) | 8 (7) |
| GB51238 | g14712_38 | fatty acid desaturase | 4 | 6 (6) | 8 (7) |
| GB51247 | g7616 | fatty acid elongase | 5 | 10 (8) | 6 (6) |
| GB51250 | g7615 | fatty acid elongase | 3 | 7 (7) | 8 (7) |
| GB53695 | g2290 | fatty amid hydrolase | 4-5* | 10-7* (9) | 6 (6) |
| GB54397 | g7610 | fatty acid elongase | 2 | 8 (NA) | 9 (NA) |
| GB54404 | g7617 | fatty acid elongase | 4 | 7 (7) | 11 (8) |

* We used both samples collected 4 days (10 bees) and 5 days (7 bees) after injection.

***Supplementary Table 11***. Oligonucleotide primers used to amplify target gene nucleotide stretches in the honey bee, required for *in situ* hybridization probe design and dsRNA synthesis*.*

| Honey beegene ID | *O. spinipes* gene ID | Forward oligonucleotide primer (5' → 3') | $T_m$ (°C) | Reverse oligonucleotide primer (5' → 3') | $T_m$ (°C) |
|---|---|---|---|---|---|
| GB40659 | g14708 | CGACTGTGGGCGCATAAAAG | 60 | TCACCCTCCTGTTGTACGGT | 60 |
| GB40681 | g7008 | TGGTCCAGTGAAGATAGCTCATAC | 64 | TTCCCATCGTATTCCCAACTGT | 60 |
| GB42218 | g14712_1 | CTCCGAAAGCAAAGAGGGAC | 60 | ACTGACGCAGATCCGCATAAC | 61 |
| GB44756 | g283 | GTGACGCAAAAGGACTTATTTGG | 61 | CTTACGCTTTACAATTACCTCTTGA | 61 |
| GB46038 | g6537 | CTCGCTGATGAACGAACGAGA | 61 | CAATGCGGCCAAACCGTAAT | 58 |
| GB48195 | g14710 | TGGCATTTCAAGATGCAGCAA | 57 | CCTTATCACCCCAACCCCAC | 63 |
| GB49380 | g7842 | GTTAGATGTGCCGCTGTTCG | 60 | TGCTGCCAGGTGATTGGATT | 60 |
| GB50627 | g2413 | TGGTCTCGGTTTGTCTTCCG | 60 | TGTAGATTTCCGACGATCTGCT | 59 |
| GB51236 | g14712_2 | TCTTCAAACAACCGCTTTCCAA | 58 | ATGCCATCCCTCACCACATC | 60 |
| GB51238 | g14712_3 | ATTATGCCATCCCTCGCCGT | 60 | TGCGGATCCTCATAATTCCCG | 61 |
| GC51247 | g7616 | ACGAGTACTTGATGAGTGGCT | 59 | AGCGAGCTTTTTCTCCGTGT | 58 |
| GB51250 | g7615 | TCACACCAGGCGGTCATTC | 59 | GCATGATCGGGTCACTCGG | 62 |
| GB52087 | g1571 | AAGCAGAGAATGTAGCTGAAGT | 58 | CGAAGACCTCTCACTCCTTGA | 61 |
| GB52590 | g3158 | CGAAAAGTTACAAGCGAACGGT | 60 | CTCGATGCACTTGACACCCT | 60 |
| GB52820 | g3059 | AGGAAATTAAGATGTCCCCGGAA | 61 | ACAGCCACAAAGAAACCTTGG | 59 |
| GB53412 | g907 | CGGTTTTTCAGGACGTTATCCT | 60 | CGTACACTCTGCGTGCATCT | 60 |
| GB53695 | g2290 | GGTCAACTCTTAACATCCCCAGT | 63 | TGTAGGCCACCAACAATCTGTAT | 61 |
| GB54302 | g7621 | GCGCTCAGGGTCGCTAGA | 61 | TGTCTTTGCTATCTCGGTCTCC | 62 |
| GB54396 | g7611 | AAAGGCCATCCAATAACGAAAGA | 59 | CCGATCGAAGCCAGCATGTA | 61 |
| GB54397 | g7610 | TCCTCGCGCAATAGAGATAACC | 62 | CGCCAAACAATGATGGACGG | 60 |
| GB54399 | g7609 | ACCTGGATTCAACTGGTGCAA | 59 | GCCACTTGACGAACCTCCTT | 60 |
| GB54401 | g7620 | GTCGGTGAACAGAAATGGCA | 59 | TTCCGGTCCCATAGCGGATA | 60 |
| GB54404 | g7617 | CGTTAAGTTCACTCCTGGTGG | 61 | CGGTTCCTAAGCCCACTGTC | 63 |
| GB55040 | g858 | TTAGACAGTAGGTTGCCGCC | 60 | CCACATACCACCTGGGACAA | 59 |
| GFP | GFP | GGAGAAGAACTTTTCACTGG | 59 | ATTCTTTTGTTTGTCTGCC | 58 |

***Supplementary Table 12***. Oligonucleotide primer sequences used to quantify the expression of target genes via quantitative reverse-transcription real-time PCR in RNAi experiments.

| Honey bee gene ID | Forward oligonucleotide primer | $T_m$ (°C) | Forward oligonucleotide primer | $T_m$ (°C) |
|---|---|---|---|---|
| GB40659 | CCGGCCAGGTTACGGATAAT | 60 | TCCTGTGGTCCCGTATCCAT | 60 |
| GB42218 | ATGGGATTTGCTGGTTCGAT | 56 | CTCTGTGGTCCCTTACCCAT | 60 |
| GB48195 | GCCGCTGTGTATGGATTGTATC | 62 | GCTGTAATTCCCAATCCAGTGC | 62 |
| GB49975 | TGCAATGTTGACAGGTTGGT | 56 | CTCTTGTCCCTTTTCTAGCTGC | 62 |
| GB51238 | GGATTAACGGAACGAAGGCA | 58 | TGTGATTCTGGTAAGAGGTTGTT | 59 |
| GB51247 | AAATCAGATCCCAGAGTGAACCA | 61 | ACGTGGAGAATAGCGTTTGGA | 59 |
| GB51250 | CAAATGGTGTTCTCAGCGTGG | 61 | ACAAAGAAGATCGTGTCCATGAAT | 60 |
| GB53695 | GGCTATCCCTCTAGCAACCAC | 63 | ACCACCAGAAGATCCACCAA | 58 |
| GB54397 | CGATCCTCGCGCAATAGAGAT | 61 | CGGAGGACAAAGAAGACCGTA | 61 |
| GB54404 | TAGCAGGATGGGGAGGTCAA | 60 | ACCACCAACAACCTTCTGCC | 60 |

**List of Supplementary Figures**

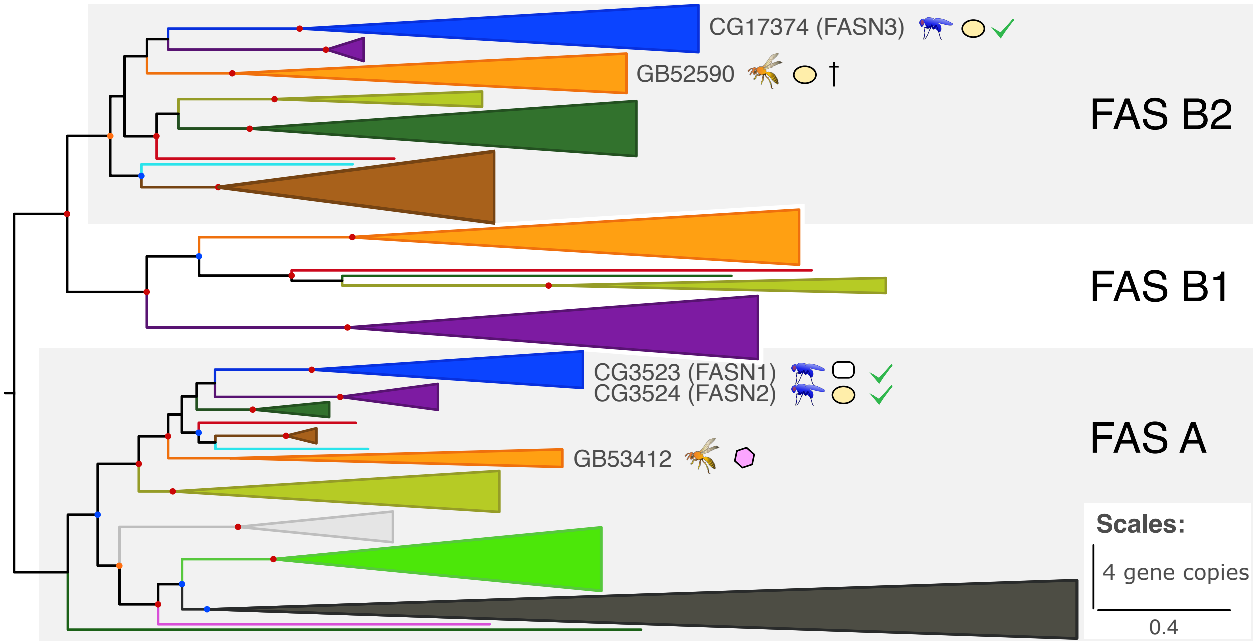


***Supplementary Figure 1***. Gene tree of fatty acid synthases from 37 Euarthropoda. Color coding and symbols are as in the legend of ***Figure 2***, the cross indicates the target gene which knockdown caused premature death in treated bees.

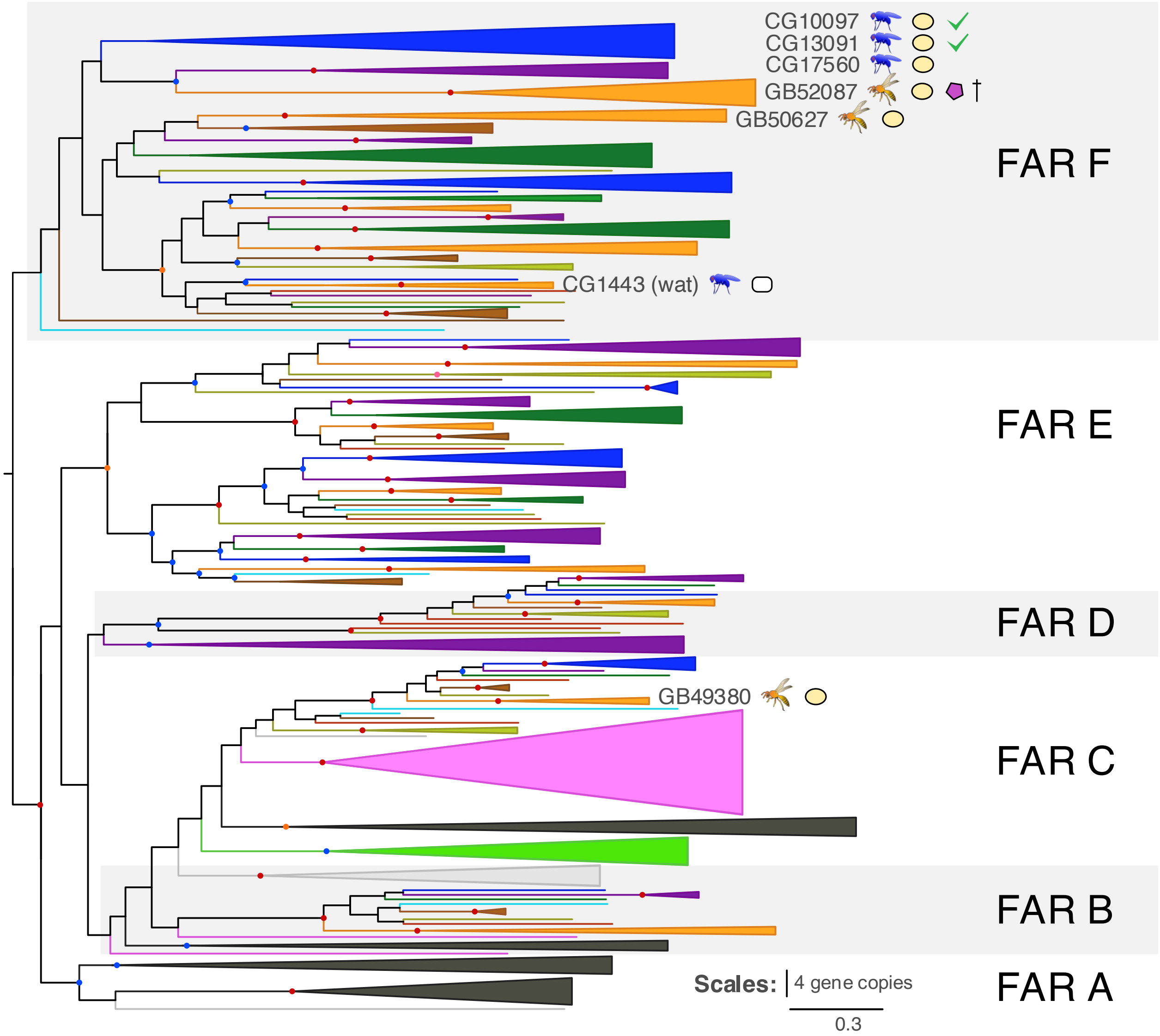


***Supplementary Figure 2***. Gene tree of fatty acyl-CoA reductases from 37 Euarthropoda. Color coding and symbols are as in the legend of ***Figure 2***, the cross indicates the target gene which knockdown caused premature death in treated bees.

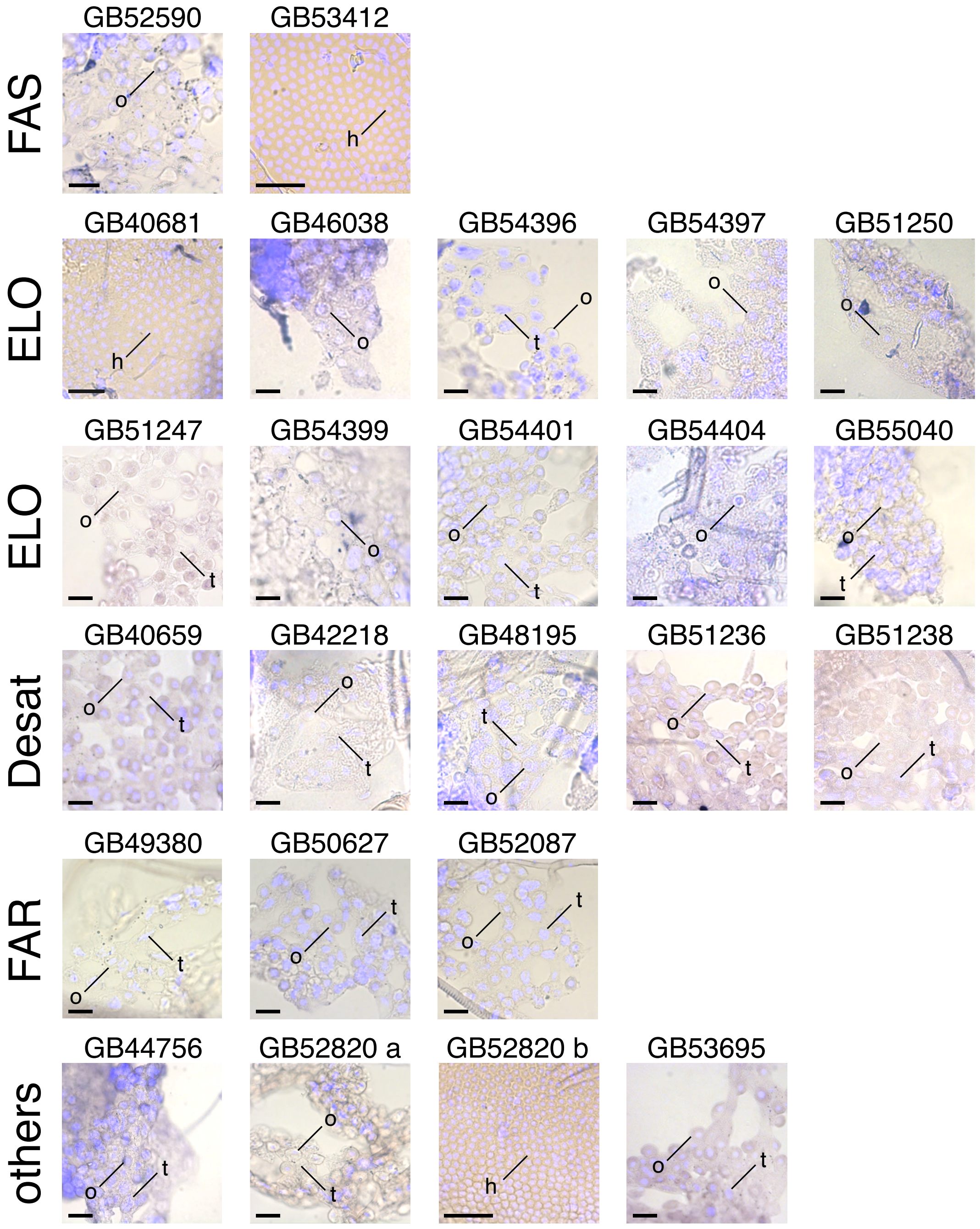


***Supplementary Figure 3***. Composition photomicrographs of worker honey bee (*Apis mellifera*) fat body cells stained with DAPI (in light blue) and RNA *in situ* hybridization experiments with the sense RNA probes (negative controls) visualizing the expression of fatty acid synthases (FAS), elongases (ELO), desaturases (Desat), fatty acid reductases (FAR), and other candidate genes (others) in three types of cells: hexagonal cells (h), oenocytes (o), and trophocytes (t). DAPI was used to counterstain the nuclei of cells. Scale bar: 50 $\mu$m.

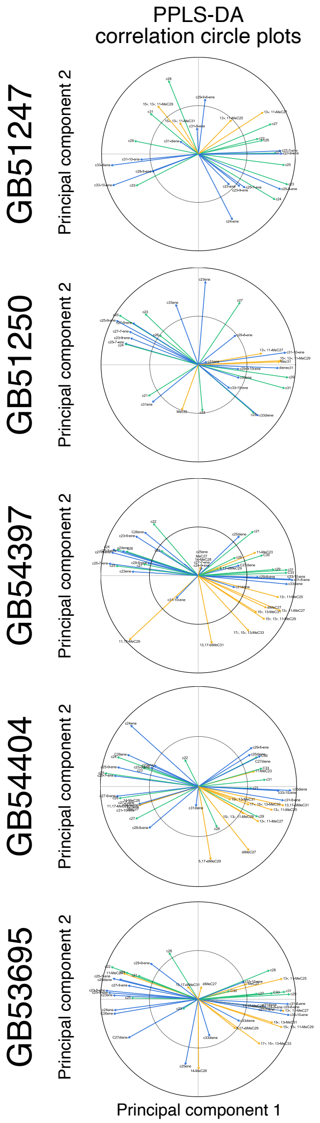


***Supplementary Figure 4****.* Effects of RNAi-mediated knockdown of four fatty acid elongases (GB51247, GB51250, GB54397, GB54404) and of one fatty acid amide hydrolase (GB53695) in worker honey bees (*Apis mellifera*) 2–5 days after dsRNA treatment. Shown are correlation circle plots from a powered partial least squares discriminant analysis (PPLS-DA) of the cuticular hydrocarbon (CHC) profile data of target gene-treated and of control bees. The correction circle plots indicate how many CHCs of a given compound class (i.e., alkanes [green], alkenes/alkadienes [blue], methyl-branched alkanes [yellow]) correlate with the first two principal components. The specific identity of the CHCs is indicated at the tip of the arrows with established acronyms. The provided information complements that given in ***Figure 5***.

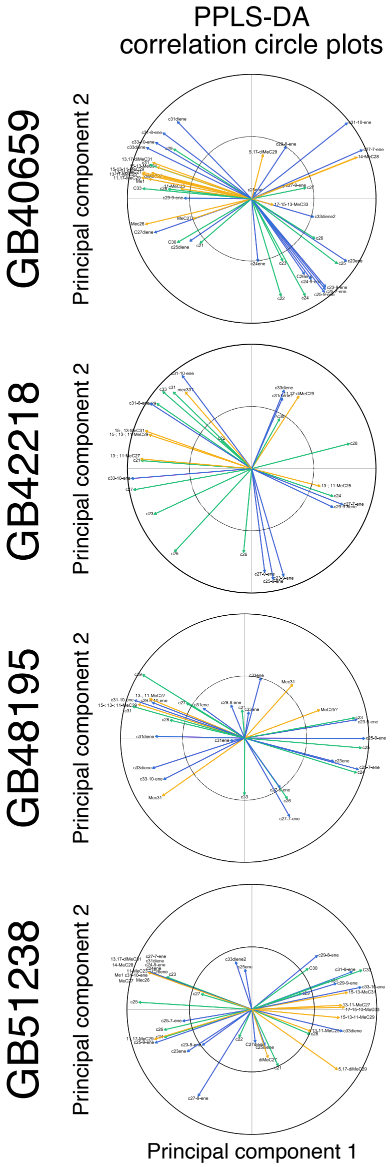


***Supplementary Figure 5***. Effects of RNAi-mediated knockdown of four fatty acid desaturases (GB40659, GB48195, GB42218, GB51238) in worker honey bees (*Apis mellifera*) 2–5 days after dsRNA treatment. Shown are correlation circle plots from a powered partial least squares discriminant analysis (PPLS-DA) of cuticular hydrocarbon (CHC) profile data of target gene-treated and of control bees. The correction circle plots indicate how many CHCs of a given compound class (i.e., alkanes [green], alkenes/alkadienes [blue], methyl-branched alkanes [yellow]) correlate with the first two principal components. The specific identity of the CHCs is indicated at the tip of the arrows with established acronyms. The provided information complements that given in ***Figure 6***.

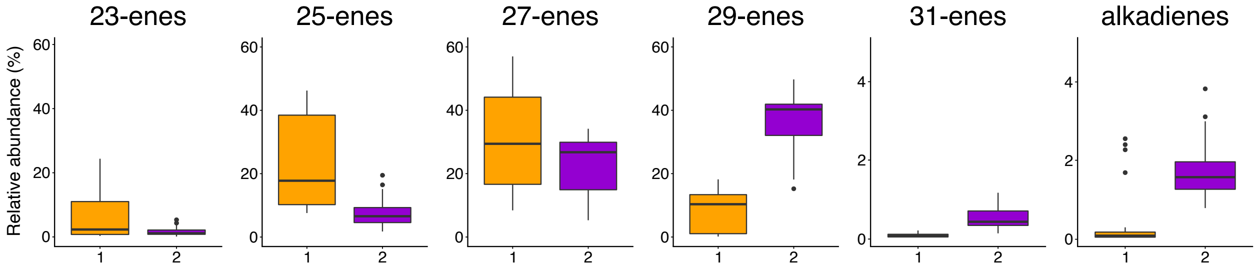


***Supplementary Figure 6***. Differences in the relative abundance of specific alkenes and of alkadienes in cuticular hydrocarbon profiles of *Odynerus spinipes* females expressing either chemotype 1 (orange) or chemotype 2 (purple).
